# A cross-kingdom interactome predicted by AlphaFold3 reveals a DNF2-centered interface required for symbiotic accommodation

**DOI:** 10.64898/2026.09.04.749384

**Authors:** Jin-Peng Gao, Feiyang Zhao, Guofeng Zhang, Qingchao Chen, Song Wu, Junfan Huang, Chun Liu, Guoliang Wang, Peng Yu, Sebastian Eves-van den Akker, Chang-Fu Tian, Thomas Ott, Jeremy D. Murray, Giles E. D. Oldroyd, Pengbo Liang, Chongjing Xia

## Abstract

Legumes convert atmospheric nitrogen into ammonium through symbiotic bacteria housed in root nodules, yet the molecular interactions between rhizobial and host proteins inside nodules remain poorly understood. Here we employed AlphaFold3 to construct a cross-kingdom interactome between *Medicago truncatula* and its symbiont *Sinorhizobium meliloti*. Screening more than 217,000 protein pairs yielded 7,137 putative interactions, providing a valuable resource for the broader symbiosis community. Within this network, we focused on DEFECTIVE IN NITROGEN FIXATION 2 (DNF2), a host protein required for rhizobial persistence within nodules. We showed that DNF2 localizes to the peribacteroid space and associates with previously uncharacterized secreted rhizobial proteins (SRPs), suggesting it may function as a hub for host-symbiont communication. Notably, knockout of two DNF2-interacting proteins, *SRP86* and *SRP485*, results in white, nitrogen-fixation-deficient nodules with abnormal symbiosomes and elevated expression of senescence-associated genes, closely phenocopying the *dnf2* loss-of-function mutant. Together, our findings define a DNF2-SRP molecular framework underlying symbiotic accommodation, and illustrate the potential of AI-guided interactome mapping to uncover molecular mechanisms of plant-microbe interactions with relevance to sustainable agriculture.

## Introduction

Cross-kingdom interactions, particularly those between plants and microorganisms, play fundamental roles in organismal fitness and survival (*1–3*). Among these interactions, legumes have evolved mutualistic relationships with nitrogen-fixing rhizobia, leading to the formation of root nodules that support carbon-nitrogen exchange (*3–5*). Within nodules, a complex yet largely unexplored cross-kingdom molecular dialogue governs the differentiation of rhizobia and their enclosure by host-derived membranes to form symbiosomes—an unusual intracellular arrangement among plant-microbe interactions.

In the model legume *Medicago truncatula*, a series of *defective in nitrogen fixation* mutants, *dnf1* to *dnf7*, have been characterized (*6, 7*). Among these, *DNF4* and *DNF7* encode Nodule-specific Cysteine-Rich (NCR) peptides that, upon processing by the *DNF1*-encoded signal peptidase complex, are translocated to the bacteroid to direct differentiation (*8–11*). In contrast, *DNF2* encodes a putative phosphatidylinositol phospholipase C-like protein implicated in bacteroid differentiation but of unknown molecular function (*12*). On the rhizobial side, apart from *nif* and *fix* genes, which encode the nitrogenase machinery and accessory proteins required for its activity, some genes involved in transport and metabolism have also been characterized (*6, 13, 14*). A recent study has applied genome-wide mariner-based transposon insertion sequencing to define hundreds of genetic regions required for effective symbiosis in *Rhizobium leguminosarum*, expanding the set of genes potentially involved in nodulation (*15*). Nevertheless, the large number of rhizobium-derived proteins have yet been functionally characterized.

Secreted proteins have emerged as key mediators of cross-kingdom communication across diverse biological interactions (*16*). Fungi and bacteria deploy extensive repertoires of secreted effector proteins to subvert host defenses and promote infection (*17–19*). Interestingly, an effector from symbiotic *Buchnera*, likely secreted via a flagellar system, is essential for colonization of aphid embryos by suppressing lysosomal activity (*20*). During rhizobia-legume symbiosis, the secretome of rhizobia is thought to play a critical role in infection and nitrogen fixation (*15, 21, 22*), yet only a few proteins have been identified (*23–25*). Functional investigation of these uncharacterized effectors will undoubtedly facilitate our understanding of endosymbiosis.

Emerging *in silico* methods, particularly AlphaFold-based structural modeling, offer a powerful approach to address this challenge by enabling systematic and unbiased identification of potential protein-protein interactions directly from sequence information (*17, 26, 27*). Yet, despite its potential, the complex cross-kingdom molecular dialogue underlying nitrogen-fixing symbiosis remains largely unexplored. In this study, we employed AlphaFold3 to screen over 217,000 *Sinorhizobium-Medicago* protein pairs, yielding an unprecedented cross-kingdom interactome resource for the broader symbiosis community. To demonstrate the utility of this resource, we focused on two previously uncharacterized secreted rhizobial proteins (SRPs), SRP86 and SRP485, which interact with DNF2 and are required for effective nitrogen fixation, potentially contributing to DNF2-mediated suppression of premature nodule senescence. Together, our study provides a powerful resource for dissecting the molecular basis of symbiotic nitrogen fixation and establishes an AI-guided framework for exploring cross-kingdom molecular interfaces.

## Results

### A candidate set: the *Sinorhizobium* secretome and host nodulation regulators

To define the molecular interface of the *Sinorhizobium-Medicago* symbiosis, we first curated a comprehensive candidate secretome for *S. meliloti* 2011 (Sm2011). Sm2011 possesses complete flagellar and Sec/Tat secretion systems and is capable of exporting large numbers of signal peptide-bearing proteins to the cell surface and into the peribacteroid space (Figure S1A). Genomic analysis identified 640 genes containing a predicted secretion peptide and without transmembrane helice, supplemented with homologs of 13 previously characterized rhizobial effectors (Table S1). These 653 proteins were considered candidate SRPs and used for subsequent analyses.

Analysis of available transcriptomic datasets for Sm2011-infected nodules (*28*) revealed that most (651 out of 653) *SRP* genes were highly expressed across 2 to 4 weeks post-inoculation (wpi) (Figure S1D, Table S1), a pattern exemplified by several candidate *SRPs* (Figure 1A), indicating their potential roles in nodulation. Predicted subcellular localization and functional annotations showed a diverse spatial distribution, including outer membrane and extracellular region (Figure S1B-S1E), reflecting the multi-stage involvement of these SRPs along nodulation process. Notably, comparison with a recent soybean nodule proteome (*22*) revealed that 215 homologous SRPs potentially localize to the symbiosome membrane and/or peribacteroid space (Figure S1F, Table S1), reinforcing our prediction regarding the localization of candidate SRPs at symbiotic interface. Thus, the rhizobial secretome presented here constitutes a transcriptionally active, spatially organized, and functionally coherent molecular toolkit deployed by symbionts throughout nodule development.

**Figure 1.**
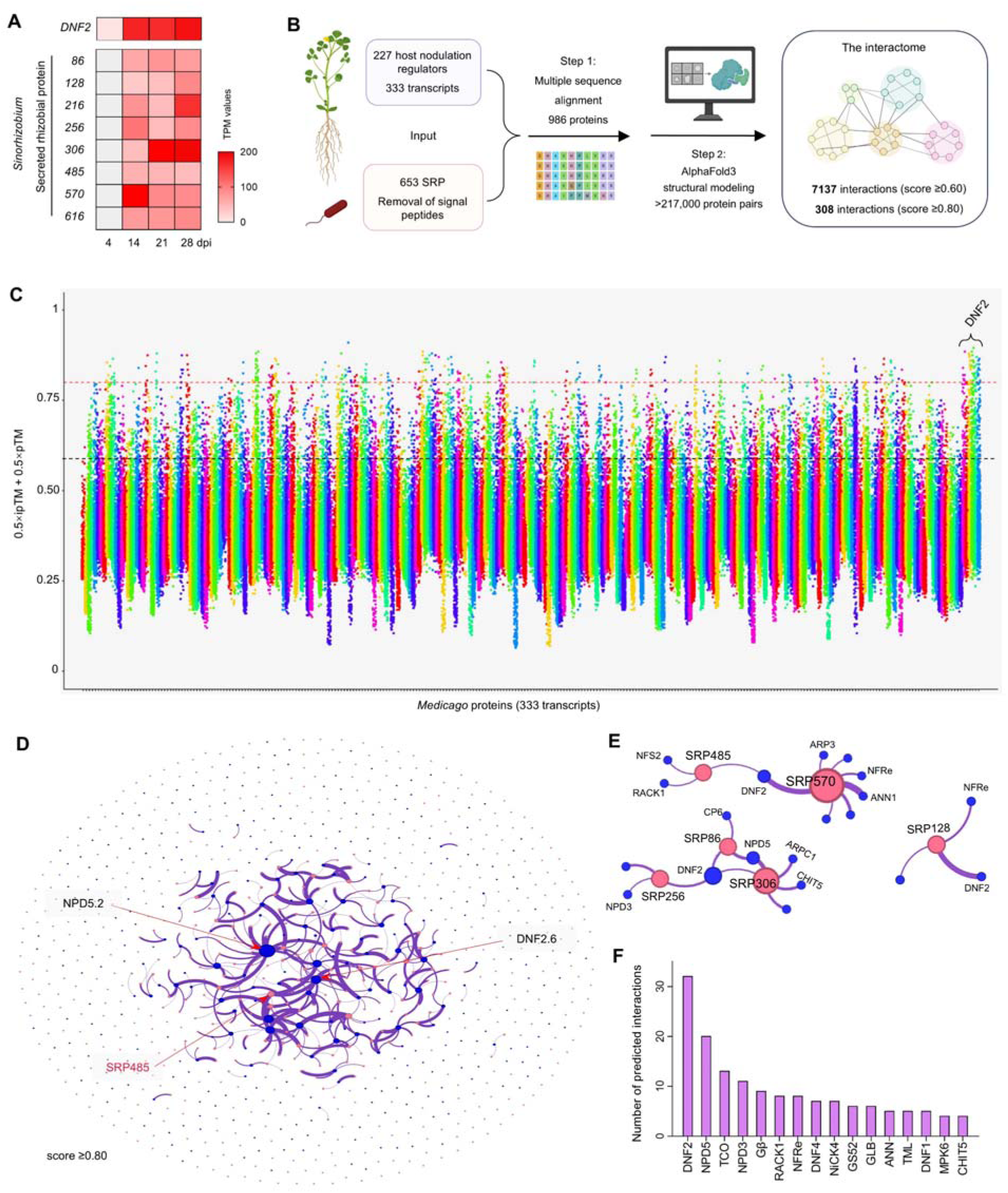
Predicted interactome of secreted rhizobial proteins and host nodulation regulators. (A) Time-course transcriptional expression analysis of candidate SRP genes at 4, 14, 21, and 28 days post inoculation (dpi). Data were reanalyzed from Sauviac et al (*28*). (B) Workflow for predicting the cross-kingdom interactome. (C) Manhattan plot of the predicted *Medicago-Rhizobium* interactome. The x-axis represents 333 *Medicago* proteins, color-coded by category, and the y-axis shows the AlphaFold3 confidence score, calculated as 0.5 × predicted template modeling (pTM) + 0.5 × interface pTM (ipTM). Red and black dashed lines indicate score thresholds of 0.8 and 0.6, respectively. (D) Network visualization of the *Medicago-Rhizobium* interactome. Blue and pink nodes represent *Medicago* and rhizobial proteins, respectively; node sizes are proportional to the number of interacting partners. Edges denote predicted interactions with scores above 0.8. (E) Network visualization of the interactions between SRPs and plant proteins. SRP570, SRP485, SRP306, SRP256, SRP128, and SRP 86 are highlighted as they interact with the higher number of *Medicago* proteins. (F) Representative plant genes that exhibited the most SRP interactions, with scores above 0.8.

Drawing on a recent review that cataloged 227 known nodulation regulators (*4*), we analyzed the expression of these genes across different nodule zones based on public transcriptome dataset (*29*) (Figure S1G). Their transcripts were differentially distributed, with zone-specific accumulation patterns evident in, for instance, the nitrogen-fixation zone (Figure S1H). To avoid omitting relevant proteins, we considered all of them as candidates and retrieved 333 corresponding *M. truncatula* proteoforms as host protein queries (Table S1). Together, these sequences served as input for cross-kingdom interactome prediction.

### AlphaFold3 modeling predicts the legume-rhizobia interactome

To evaluate the utility of AlphaFold3 for predicting protein complexes involved in nitrogen-fixing symbiosis, we followed a composite confidence metric defined as 0.5 × predicted template modeling score (pTM) + 0.5 × interface pTM (ipTM), to rank predictions. This metric integrates confidence in the overall complex structure and the predicted interaction interface (*27*). We began by assessing a set of well-characterized protein interactions that are essential for nodulation. The predicted model of the Nod factor receptor NFR1-NFR5 complex and the NODULATION SIGNALING PATHWAY 1 (NSP1)-NSP2 heterodimer agreed with experimental data (*30–32*), and achieved high confidence scores of >0.80 (Figure S2A and S2B). The interaction between the effector Nodulation Outer Protein NopM and NFR5 (*33*), was predicted with high confidence (score =0.78, Figure S2C).

Building on previously reported benchmarks and our current results, we conclude that AlphaFold3 possesses reasonable capability to predict nodulation-related protein-protein interactions. We therefore subjected 653 rhizobial and 333 plant protein sequences to a structural modeling pipeline (Figure 1B), in which all possible binary combinations were folded using AlphaFold3 to generate over 217,000 predicted protein-protein complexes.

We first applied a moderate-confidence threshold of ≥0.60 to maximize sensitivity, identifying 7,137 putative interactions (Figure 1C, Figure S3A, Table S2). We further applied a more stringent screening threshold (score ≥0.80), which yielded 308 high confidence protein-protein interaction pairs (Figure 1C, Table S2). At this stringent cutoff, a total of 189 SRPs were each predicted to interact with at least one host protein, collectively targeting 68 *M. truncatula* genes, corresponding to 89 variants (Figure 1D, Figure S3B). Network reconstruction of this predicted interactome revealed a highly non-uniform topology, characterized by specific host regulators serving as central hubs rather than a dispersed web of interactions (Figure 1D, Figure S3A).

From the rhizobial side, SRP485, a putative phosphonate ABC transporter substrate-binding protein, emerged as a multi-target effector (Figure 1D and 1E, Figure S3C). Specifically, it was predicted to interact with DNF2 (*12*), NFS2, an NCR peptide governing symbiotic compatibility (*34*), and the scaffold protein RACK1 (Receptor for activated C kinase) (*35*). In addition, SRP256, a copper-binding protein, was predicted to interact with DNF2 and the nodule-specific PLAT domain protein NPD3 (*36*) (Figure 1E). Another predicted effector, SRP86, a Domain of Unknown Function 2155 protein, was predicted to associate with DNF2, NPD5, and CP6 (a cysteine protease linked to nodule senescence) (*37*). Moreover, SRP306 (a YceI family protein), was predicted to interact with NPD5, ARPC1 (a subunit of the ARP2/3 complex) (*38*), and CHIT5 (a chitinase) (*39*). SRP570, a SnoaL-like domain-containing protein, was predicted to interact with Nod factor receptor NFRe (*40*), Actin-Related Protein 3 (ARP3) (*41*), and Annexin 1 (ANN1) (*42*) (Figure 1E, Figure S3C).

On the plant side, the predicted cross-kingdom interactions spanned nearly all stages of nodulation. In signal perception and early recognition, the chitinase CHIT5, which can cleave Nod factors (*39*), was predicted to interact with 4 SRPs, while the receptor NFRe was linked to 8 SRPs (Figure 1F). The heterotrimeric Gβ protein (*43*) and the scaffold protein RACK1 were predicted to interact with 9 and 8 SRPs, respectively. During rhizobial infection and organogenesis, ANN1, a regulator of infection thread formation, was targeted by 5 SRPs, alongside TRICOT, a carboxypeptidase that regulates infection and nodule organogenesis (*44*), which showed 13 predicted interactions. For systemic regulation, Too Much Love (TML), a known synergistic modulator of nodulation (*45*), was predicted to be targeted by 5 SRPs. Finally, in nitrogen fixation, DNF2 ranked first with 32 predicted interactions, followed by NPD5 with 20 (Figure 1F, Figure S3E, S3F), while the hemoglobin GLB1 (*46*) showed 6 predicted interactions.

Together, we define this network as the legume-rhizobia interactome, and provide an interactive web database (http://20.77.1.79/loadcsv.html) for the community to browse and validate the uncharted symbiotic interfaces (Figure S3G).

### DNF2 physically interacts with multiple novel SRPs

To demonstrate the utility of this resource, we focused on DNF2, a host protein that not only showed the most abundant interactions with SRPs (Figure 1F), but also exhibited high expression in the infection and nitrogen-fixation zones of nodules (Figure S1H).

To validate the identified interactions, we employed a split-luciferase (split-LUC) complementation assay. Constructs were generated to fuse the DNF2 to N-terminal LUC, and each of the 10 SRPs to the C-terminal fragment. These construct pairs were co-expressed in *Nicotiana benthamiana* leaves. The assay showed at least seven interaction partners of DNF2 among the 10 tested, with particularly strong signals observed for pairs involving SRP86 and SRP485 (Figure 2B and 2C, Figure S4). The interactions between DNF2 and these two SRPs were further confirmed by co-immunoprecipitation *in vivo* (Figure 2D and 2E).

**Figure 2.**
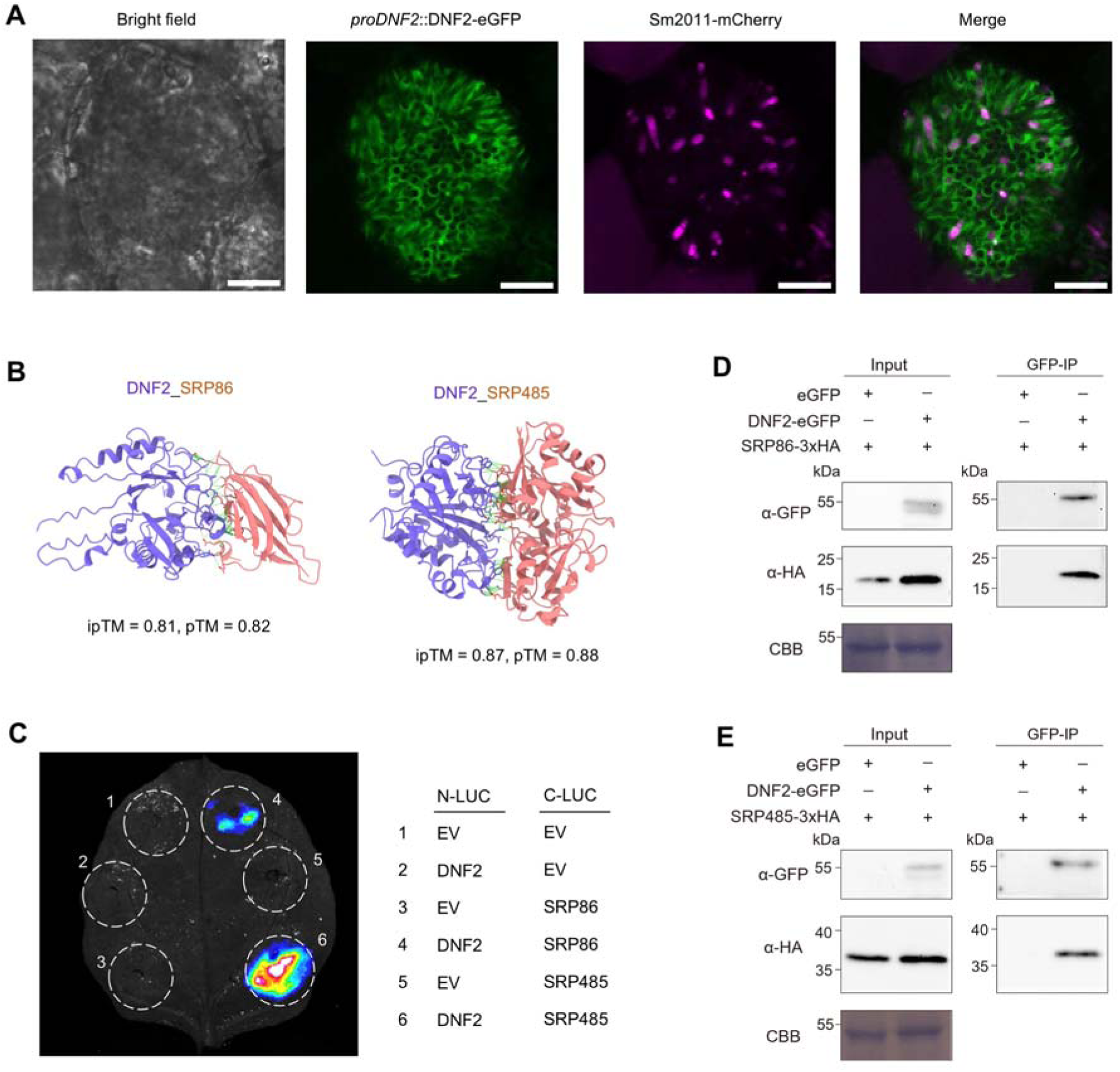
DNF2 interacts with multiple rhizobial proteins. (A) Confocal images of *M. truncatula* nodule cells showing subcellular localization of DNF2-eGFP and rhizobia Sm2011-mCherry. Scale bars = 10 μm. (B) Predicted interaction models of DNF2 with SRP86 (left) and SRP485 (right) generated by AlphaFold 3. The predicted scores are indicated in the figure. (C) Split luciferase (LUC) complementation assays between DNF2.6 and SRPs. The N-terminal fragment of LUC (nLUC)-tagged DNF2 was co-infiltrated into *N. benthamiana* leaves together with the C-terminal fragment of LUC (cLUC)-tagged SRP86 or SRP485. (**D**, **E**) Co-immunoprecipitation (co-IP) assays of SRP86-HA or SRP485-HA and DNF2-eGFP from total proteins extracted from *N. benthamiana* leaves. Proteins were immunoprecipitated (IP) with anti-GFP beads and analyzed by western blotting. CBB (Coomassie brilliant blue) staining showed equal total protein loading.

We further showed DNF2-eGFP, driven by the native promoter, was exclusively detected in infected cells of nodules and localized to the peribacteroid space, where it surrounded differentiating rhizobia (Figure 2A). Together, this distinct spatial localization pattern strongly supports the direct involvement of DNF2 at the host-symbiont interface, positioning it as a potential hub for interacting with SRPs within the symbiotic network.

### DNF2-interacting SRPs are required for nitrogen fixation in nodules

To investigate the roles of these DNF2-associating SRPs in the nitrogen-fixing symbiosis, we generated a series of deletion mutants in Sm2011, including *srp86*, *srp256*, and *srp485* (Figure S5). Most *M. truncatula* wild-type A17 or R108 plants inoculated with the rhizobial single deletion mutants displayed slightly but significantly reduced shoot length and increased number of white nodules (Figure 3A-D, Figure S6). However, the majority of nodules induced by these rhizobial mutants remained pink and elongated, comparable to those induced by wild-type Sm2011, indicating that active nitrogen fixation was largely maintained (Figure 3C). Given that DNF2 interacts with all three SRPs, we hypothesized functional redundancy or alternative mechanisms among them.

**Figure 3.**
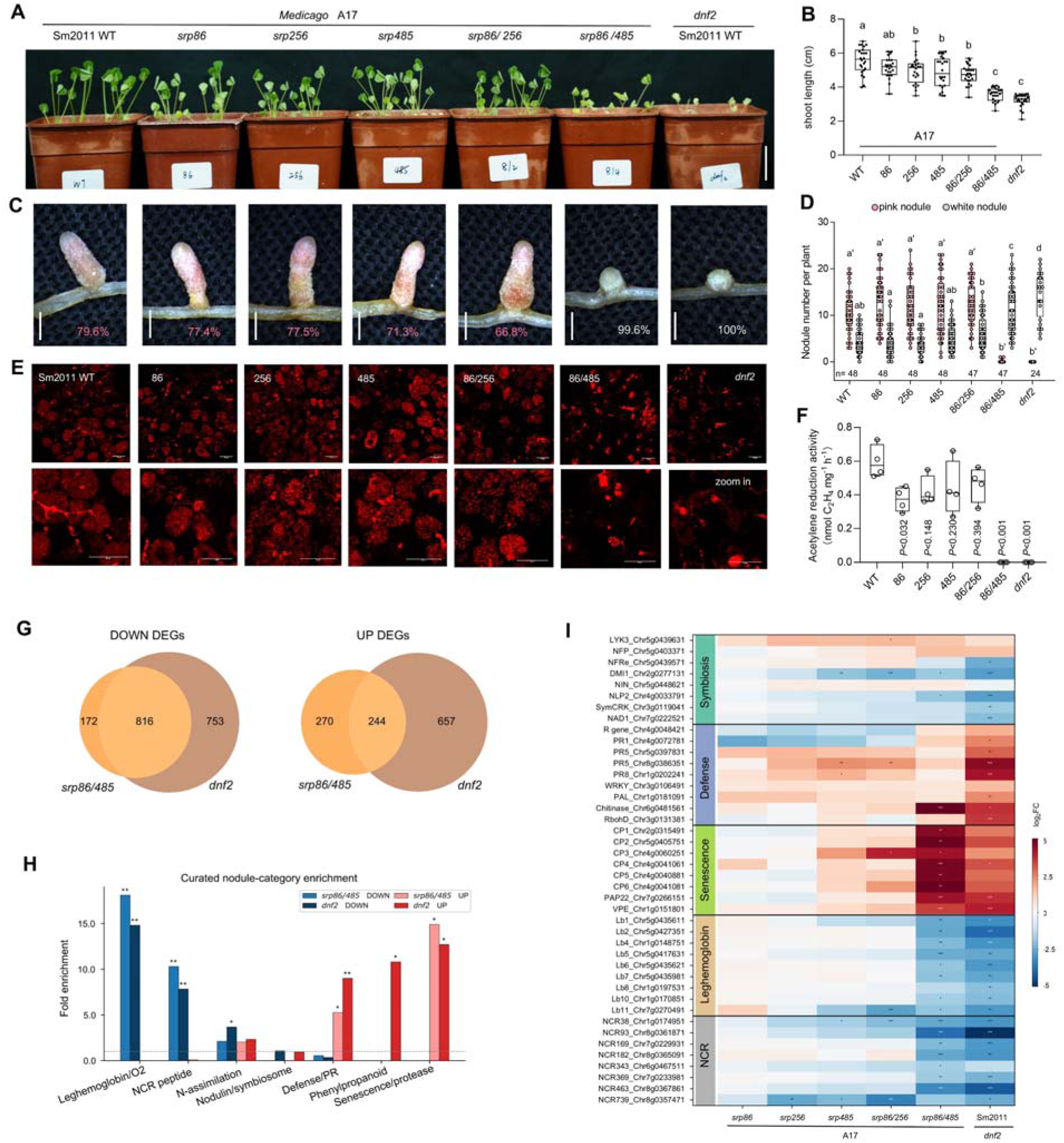
Secreted rhizobial proteins are essential for symbiotic nitrogen fixation. (A) Growth phenotype of *Medicago truncatula* plants at 3 weeks post-inoculation (wpi) with the wild-type strain *S. meliloti* 2011 (Sm2011, WT), or the *srp* mutants. Representative images showing overall plant development. Scale bars = 3 cm. (B) Quantification of shoot length at 3 wpi. Boxes show the first quartile, median, and third quartile; whiskers show minimum and maximum values; dots show data points (n = 24). Statistical significance was determined using one-way ANOVA followed by Tukey’s HSD test, different letters denote significant differences. (C) Representative stereomicroscopy images of mature nodules harvested at 3 wpi. Scale bars = 1 mm. Red and white numbers represent the percentages of pink and white nodules, respectively. (D) Quantification of the proportion of pink (functional) versus white (non-functional) nodules at 3 wpi. Samples size (n) was indicated on the figure. (E) Representative confocal microscopy images of nodule nitrogen fixation zones. Red fluorescence indicates *S. meliloti* Sm2011 expressing mCherry (Sm2011-mCherry). Scale bars = 50 µm. (F) Acetylene reduction assay (ARA) measuring nitrogenase activity in nodules formed by the indicated strains. Statistical significance was assessed using a two-tailed Student’s *t*-test. ARA activity is expressed as ethylene production per mg of fresh nodule weight per hour (nmol C₂H₄ mg⁻¹ FW h⁻¹). (G) Overlap of differentially expressed genes (DEGs) between *srp86/485* and the host *dnf2* mutant (down– and up-regulated). (H) Curated nodule-category enrichment for the four directional sets. Symbiotic categories (leghemoglobin/O₂, NCR, N-assimilation) are over-represented among down-regulated genes of both genotypes; defense/PR and phenylpropanoid are over-represented only among *dnf2* up-regulated genes. (I) Fold-change of expression of representative genes in nodules at 3 wpi.

To further explore this possibility, we generated *srp86/srp256* and *srp86/srp485* double mutants (Figure S5). Interestingly, *srp86/srp485*, but not *srp86/srp256*, induced predominantly small, round and white nodules at 3 and 4 wpi in both A17 and R108, whereas the wild-type Sm2011 induced elongated pink nodules (Figure 3C, Figure S6). Consistent with this, plants infected by *srp86/srp485* double mutant strain exhibited typical nitrogen-deficient phenotype, as revealed by yellowish leaves and significantly reduced aerial biomass (Figure 3A, Figure S6). Confocal microscopy analysis of nodule sections revealed aberrant symbiosome development in plants inoculated with the double mutant strain (Figure 3E). Consistently, acetylene reduction assays confirmed that nitrogenase activity was significantly reduced in these nodules induced by the *srp86/srp485* mutant strain (Figure 3F). Together, these results demonstrate that *SRP86* and *SRP485* are critical for symbiotic nitrogen fixation, with their nodulation phenotypes mirroring that previously described for the *dnf2* mutant (*7, 12*), thus suggesting that DNF2 and these SRPs form a functional module.

### The DNF2-SRP module maintains symbiotic gene expression and prevents nodule senescence

To further elucidate the mechanisms underlying the functional relationship of DNF2 and SRPs, we performed bulk RNA-seq analysis on nodules infected by *srp* single and double mutants, as well as *dnf2* nodules (Figure S7). The single mutants of *srp86* and *srp485* showed a moderate response, whereas the *srp86/srp485* double mutant produced 1,502 differentially expressed genes, with down-regulation dominating (988 down, 514 up). This down-regulation was overwhelmingly emergent: 84% of the down-regulated genes were altered in neither single mutant, indicating a synergistic, not additive, collapse (Figure S7D). Compared with *dnf2*, the *srp86/srp485* mutant shared a large core of 816 down-regulated genes and 244 up-regulated genes (Figure 3G). Overall, the transcriptional profiles revealed a transcriptional collapse that selectively phenocopied the host *dnf2* mutant.

Interestingly, the shared down-regulated set converged on the symbiotic machinery—leghemoglobins, oxygen transport, NCR peptides, and nitrogen assimilation—showing that the double mutant faithfully recapitulated the symbiotic arm of *dnf2* (Figure 3H). In contrast, the up-regulated arms diverged (Figure 3H). *dnf2* mounted an acute immune response enriched for defense signaling, pathogenesis-related genes, and phytoalexin biosynthesis, whereas the *srp* mutants induced a more modest immune response; its up-regulated genes instead reflected proteolysis and senescence-like processes (Figure 3H and 3I). Quantitatively, the *srp86/srp485* mutant recapitulated 52% of *dnf2*-down-regulated symbiotic genes but only 27% of the up-regulated defense genes (Figure S7I). Thus, loss of bacterial *SRP86/SRP485* disables the DNF2-dependent symbiotic accommodation programme, causing a senescence-prone nodule, while the host DNF2 protein still suppresses immunity independently of these effectors. The DNF2-SRP module therefore sustains symbiotic gene expression and prevents nodule senescence, whereas defense suppression is an intrinsic function of DNF2.

### The DNF2-SRP module appears to be conserved across rhizobia-legume interactions

To test whether the DNF2-SRP module is conserved in other rhizobia-legume interactions, we first investigated whether *DNF2* is conserved in the legumes. Phylogenomic analysis indicated that DNF2 is legume-specific and structural predictions for DNF2 orthologs from *Medicago*, pea (*Pisum sativum*), and soybean (*Glycine max*) revealed a high degree of structural conservation, so likely DNF2 has a conserved function in nodules (Figure 4A and 4B). A *DNF2*-like gene, from which *DNF2* may have evolved, is conserved in most plants and exhibits ubiquitous expression including in non-inoculated roots (Figure S1I), a pattern distinctly different from that of *DNF2*.

**Figure 4.**
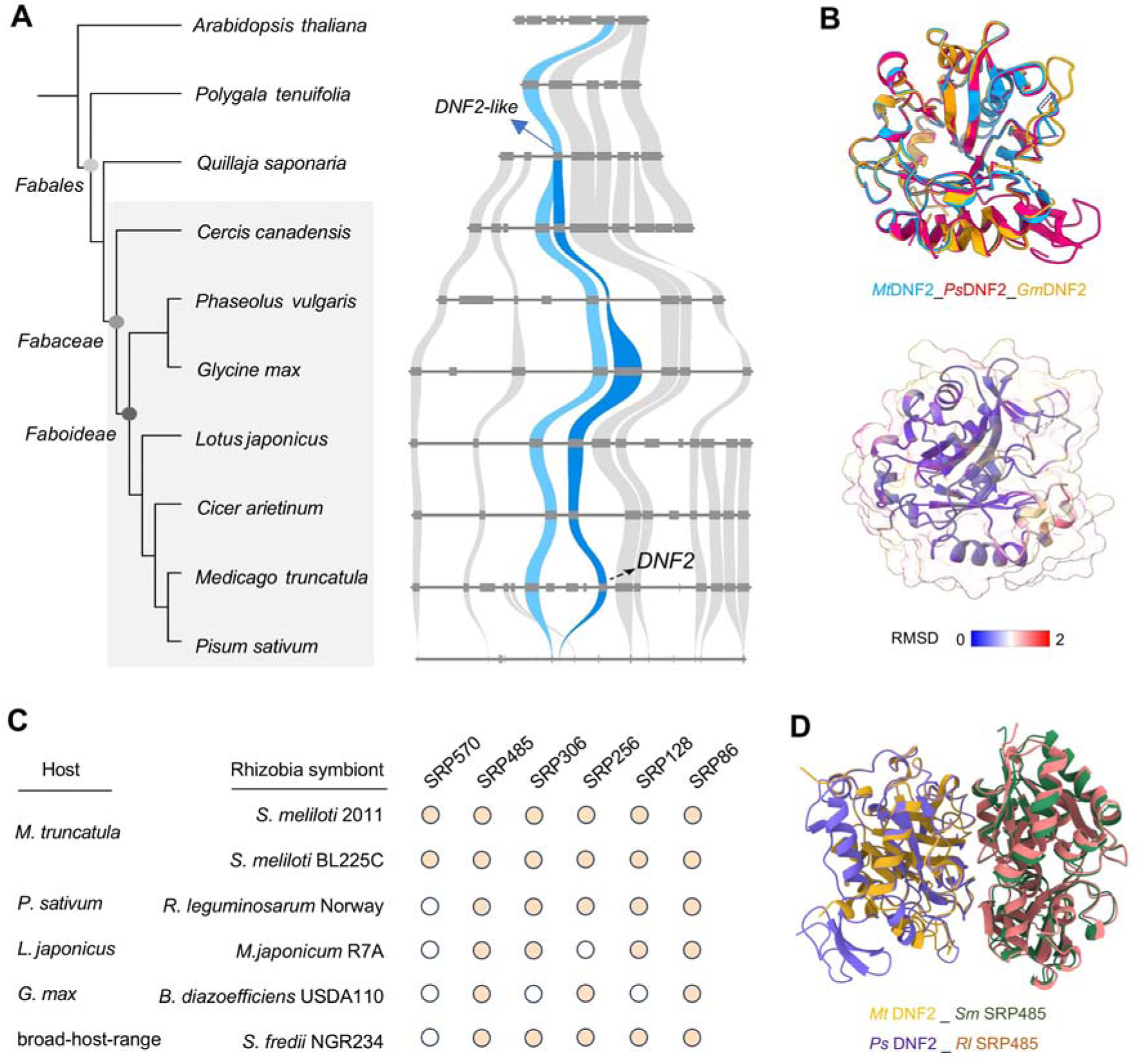
The DNF2-SRP module appears to be conserved in legume-rhizobia symbiosis. (A) Synteny analysis of DNF2 in representative non-Leguminosae and Leguminosae species. Phylogenetic tree of leguminous and non-leguminous plants (Left). Homologous genes within each syntenic block are connected by color-coded lines. (B) Structural overlay of DNF2 proteins from *Medicago*, pea (*Pisum sativum*), and soybean (*Glycine max*). The root mean square deviation (RMSD) values are indicated to reflect the structural similarity among the homologs. (C) Distribution of candidate SRP homologs across representative rhizobial strains. Cream-colored circles indicate the presence of homologous proteins, whereas white circles indicate their absence. (D) Predicted structures of *Medicago* DNF2 and pea (*Pisum sativum*) DNF2 in complex with their respective rhizobial SRP485 counterparts, generated by AlphaFold 3.

Interestingly, several SRPs, including SRP86 and SRP485, have orthologs in other rhizobia (Figure 4C and Figure S8). AlphaFold3 predicts that DNF2 interacts with high confidence with SRP485 from other rhizobia, such as those in *Rhizobium leguminosarum*, a symbiont of pea (Figure 4D).

Collectively, as we have demonstrated in the *Medicago-Sinorhizobium* association, the DNF2-SRP interaction is likely conserved in other legume-rhizobia interactions, indicating its crucial function for the establishment of nitrogen-fixing symbiosis.

## Discussion

The host-microbe interface is governed by a complex exchange of secreted signals, including effector proteins (*17, 20, 47*). By systematically mapping the *Medicago-Sinorhizobium* interactome using AlphaFold3, we identified the host phospholipase-like protein DNF2 as an interkingdom hub engaging multiple previously uncharacterized SRPs. While DNF2 has been linked to defense suppression via cleavage of immune receptors LysM receptor-like proteins LYM1/LYM2 (*48, 49*), our data indicate that DNF2 mediated regulation may involve layered functional aspects. Unlike *dnf2* mutants which triggers strong immunity, the loss of *SRP86*/*SRP485* leads to a strong defect in symbiotic maintenance with a comparatively muted immune response (Figure 3G-I). This indicates that engagement of DNF2 by bacterial SRP86 and SRP485 is essential primarily for sustaining the symbiotic accommodation program, which maintains leghemoglobin, NCR peptides, and active nitrogen fixation. Furthermore, emerging evidence has shown that phosphatidylinositol 4,5-bisphosphate [PI(4,5)P_2_] is enriched at symbiotic interfaces during both arbuscular mycorrhizal and rhizobial colonization, but not at those associated with the pathogenic oomycete *Phytophthora palmivora* (*50, 51*). It is plausible that DNF2-SRPs regulate the local accumulation and distribution of these lipid signatures.

Our findings, supported by previous studies (*15, 22*), underscore that rhizobia deploy a vast, transcriptionally active secretome to negotiate endosymbiosis. In a parallel study, AlphaFold2 predicted structures of Type III Secretion System (T3SS) effectors from rhizobia and pathogens, which presumably mimic DNA-binding folds in the nucleus (*25*). The present work, however, centers on the broader repertoire of secreted proteins that operate beyond the T3SS to engage the plant host. Indeed, AlphaFold-Multimer was recently used to predict the interactions between secreted proteins and defense-related hydrolases at the tomato-pathogen interface (*17*). This perspective is reinforced by a recent report on the symbiotic bacterium *Buchnera,* demonstrating that a putatively flagellar-secreted protein translocates into aphid cells and is essential for colonization (*20*).

Although core SRP and DNF2 interactions appear structurally conserved across diverse legume and rhizobia pairings (Figure 4), whether strain specific SRP repertoires influence host selectivity—or whether host processing machinery like DNF1 and NCR peptides directly interface with rhizobial effectors—remains an intriguing open question.

Traditional validation of effector function is low-throughput and hindered by sequence diversity and rapid evolution (*52*). We offer a complementary strategy for uncharacterized effectors across diverse host and microbe systems. One limitation of *in silico* prediction is the potential for false positives, which may arise when candidate proteins do not co-localize within the same cellular compartment or are expressed asynchronously *in planta*. Conversely, false negatives could occur due to proteins with intrinsically disordered regions or poor sequence alignment depth.

Collectively, our findings define a molecular framework for DNF2-SRP function in symbiotic accommodation and, more broadly, provide a strategy for dissecting uncharted interkingdom interfaces.

## Methods

### Plant materials and growth conditions

*Medicago truncatula* ecotype A17 and R108, as well as *dnf2* mutant (A17 background) (*7*) were used in this study. *Nicotiana benthamiana* was used for protein-protein interaction analyses. All plants were grown in controlled environment chambers with 16 h of light, 8 h of dark at 22°C with 55% humidity, and the light intensity was 200 μmol m^-2^ s^-1^.

### Microbial strains

The *Sinorhizobium meliloti* 2011 (Sm2011) was used in this study for nodulation assays. *Escherichia coli* DH5α was used for gene cloning. *Agrobacterium rhizogenes* ARqua1 strain was used for *M. truncatula* hairy root transformation. The strain *A. tumefaciens* GV3101 was used for transient transformation in *N. benthamiana*.

### Characterization of secreted rhizobial proteins

To predict putative effectors of Sm2011, proteome was retrieved from NCBI (RefSeq assembly: GCF_000346065.1). For each protein, the presence and location of signal peptide cleavage sites were predicted using SignalP6 (*53*). The presence of transmembrane helices was predicted using DeepTHMM (v1.0.24) (*54*). A total 640 proteins containing signal peptides and lacking transmembrane helices were considered as putative effectors. In addition, 13 previously described effectors (NopA, NopB, NopC, NopD, NopE, NopI, NopL, NopM, NopP, NopT, InnB, ErnA and Bel2-5) were also used (Table S1).

Functional characterization of the secretome was performed through multi-layered annotation. Comprehensive domain architecture and Gene Ontology (GO) term assignments were generated using InterProScan 6.0.0 (*55*) against integrated databases (Pfam, SMART, PANTHER, SUPERFAMILY), with GO terms propagated to enable enrichment analysis. Subcellular localization was predicted using DeepLocPro 1.0.0 (*56*) to assess targeting beyond the bacterial membrane. Metabolic pathway context was established through KEGG orthology assignment via KofamScan 1.3.0 (*57*) using prokaryote-specific HMM profiles (prokaryote.hal). GO and KEGG enrichment analyses were conducted by comparing the secreted protein subset against the complete *S. meliloti* proteome background.

### Transcriptomic analysis of the rhizobial secretome in nodules

To assess the functional relevance of computationally predicted secreted effectors, we leveraged publicly available RNA-seq datasets (SRA accessions SRR18299090-SRR18299092, SRR18299142-SRR18299149) derived from *S. meliloti* 2011 isolated from *M. truncatula* nodules at multiple developmental stages (2, 3, and 4 weeks post-inoculation) (*28*). Raw paired-end reads (Illumina) underwent rigorous quality control: adapter sequences and low-quality bases (Phred score < 20) were trimmed using Trimmomatic v0.39 (*58*) with sliding window (4:20) and minimum length (60 bp) parameters, retaining only properly paired reads for downstream analysis. High-quality reads were aligned to the *S. meliloti* 2011 reference genome (GCF_000346065.1) using STAR v2.7.10b (*59*) with stringent settings (-outFilterMultimapNmax 1) to ensure unambiguous mapping of reads to protein-coding loci. Transcript abundances were quantified in transcripts per million (TPM) and fragments per kilobase per million (FPKM) using StringTie v3.0.0 (*60*), with expression estimates normalized for gene length and sequencing depth to enable cross-sample comparability. Expression profiles of the predicted secreted proteins were extracted from the genome-wide TPM matrix and subjected to clustering using ClusterGVis (*61*).

### Prediction of known interactions using AlphaFold3

For the known interactions, LjNFR1-LjNFR5 (*30*), MtNSP1-MtNSP2 (*32*), and NopM-LjNFR5 (*33*) complexes were modeled in two modes, the full length and the functional domains, using AlphaFold3 (v3.1) (*27*). For NopM, the catalytically inactive mutant NopMC338A (NCBI accession AAB91674) was used to capture stable protein complex. To capture the most accurate model, 50 seeds were set for AlphaFold3 inference.

### Prediction of the *Medicago-Rhizobium* interactome

Secreted rhizobial proteins after removing signal peptides were used to predict interactions with the 227 known nodulation regulators identified previously (*4*). To predict protein complex structures at scale using Cambridge University’s Research Computing Services (https://www.hpc.cam.ac.uk/), we implemented a high-throughput AlphaFold3 pipeline. The workflow was separated into two phases: first, evolutionary features and MSAs were generated individually for each protein using 8 CPUs. Second, these precomputed features were merged using custom scripts for each protein pair and integrated into complex-specific inputs for GPU-accelerated inference. Total of 10 models were generated for each complex by AlphaFold3 default setting (one random seed with 10 samples). At this stage one GPU (NVIDIA A100-SXM-80GB) was used for each protein pair. The entire screen consumed a total of >10872 Graphics Processing Unit (GPU) hours (>1.25 GPU years).

### Vector construction

All gene and promoter sequences used in this study were synthesized (GeneArt, Invitrogen, Thermo Fisher Scientific, Waltham, MA, USA). For subcellular localization of DNF2, the putative 4 kb promoter region and the coding sequence of *DNF2* were synthesized with appropriate overhangs for Golden Gate assembly, following silent mutagenesis to remove internal *BsaI* and *BpiI* restriction sites. For hairy root transformation, the assemblies of level 1 and level 2 plasmids were using Golden Gate cloning system. All Golden Gate functional modules are available through the ENSA project (https://www.ensa.ac.uk/).

### Spilt luciferase complementation assay

*A. tumefaciens* strain GV3101 harboring the plasmid of interest was cultured overnight at 28°C in LB medium. Bacteria were collected and re-suspended in infiltration buffer (10 mM MgCl_2_, 10 mM MES of pH 5.7, 200 µM acetosyringone), adjusted to a final density of OD_600_=0.3 and incubated at room temperature for 2 h with gentle agitation before being infiltrated into the *N. benthamiana* leaves. After 2-3 days of infiltration, the leaves were collected and sprayed with a solution of D-Luciferin potassium salt (E1500, Promega, Madison, WI, USA). The images were acquired using a ImageQuant 800 system (GE Healthcare, Chicago, IL, USA).

### Co-Immunoprecipitation

*A. tumefaciens* strain GV3101 carrying the indicated constructs was infiltrated into *N. benthamiana* leaves. Three days after infiltration, leaves were harvested frozen in liquid nitrogen, and homogenized in 1 mL of cold extraction buffer [50 mM Tris/HCl of pH 7.5, 100 mM NaCl, 0.5% Triton X-100, 10 mM β-mercaptoethanol and Protease Inhibitor Cocktail (Roche, Basel, Switzerland)]. Crude extracts were then centrifuged at 16000 g for 10 minutes at 4°C, and the supernatant was used to determine protein concentrations by Bradford assay (Bio-Rad, Hercules, CA, USA). For co-immunoprecipitation, 25 μl of GFP-Trap magnetic agarose (Chromotek, Martinsried, Germany) was added into each supernatant and incubated for two hours at 4°C with constant agitation. Beads were separated using magnetic rack and washed three times with extraction buffer. Samples were eluted in Laemmli buffer at 95°C for 5 minutes, and loaded on 12% SDS-PAGE gels at 90-130 V for 2 hours. The proteins were then transferred onto a PVDF membrane (0.2 µm pore size) using Trans-blot Turbo system (BioRad, Hercules, CA, USA). Membranes were blocked in 5% (w/v) milk-TBST for 1 hour, and incubated with anti-GFP (1:5000, Cat# 632381, Takara, Kusatsu, Japan) or anti-HA (1:4000, Cat# 12013819001, Roche, Basel, Switzerland) antibody at 4°C overnight. Membranes were washed four times with TBST for 30 minutes in total. Where required, peroxidase-conjugated anti-mouse (1:5000, Cat# A4416, Sigma-Aldrich, St. Louis, MO, USA) was applied for 2 hours at room temperature. Protein signals were detected using Clarity Western ECL Substrate (Bio-Rad, Hercules, CA, USA) and visualized with an ECL ChemoCam Imager (Intas Science Imaging, Germany).

### Targeted gene knockout in *Rhizobium* via tri-parental mating

The target gene deletion mutant was using homologous recombination based on the pJQ200SK plasmid combined with sacB-mediated sucrose counter-selection. The linearized vector and PCR-amplified upstream/downstream flanking fragments of the target gene were assembled via seamless cloning and transferred into the recipient rhizobium via tri-parental conjugation. Single-crossover integrants, selected on Gent/Tet plates, were verified by PCR. Subsequent non selective cultivation was performed on a tetracycline-resistant medium to facilitate the second exchange, thereby promoting a second recombination event. Double crossover mutants were selected on sucrose containing tetracycline plates. Final mutants were confirmed by PCR and sequencing of the deletion region. The primers used are list in the Table S4.

### Acetylene reduction assay

Fresh nodulated roots from three plants were surface-dried, placed in 100 mL sealed vials, and 10 mL of air was replaced with 10 mL of acetylene. After 2 h incubation at 28°C, 2 mL of reaction gas was drawn and injected into a 10 mL liquid-sealed quenching bottle. Subsequently, 100 μL of headspace gas was injected into a gas chromatograph (7820A, Agilent Technology, Santa Clara, CA, USA) to measure ethylene production. The ethylene peak area was recorded after 1.5 min and three technical replicates were performed. Nodules were then excised, weighed, and acetylene reduction assay was calculated as nmol ethylene per mg fresh weight per h.

### Hairy root transformation

*M. truncatula* seedlings were transformed via *A. rhizogenes* strain ARqua1 mediated hairy root transformation. Seed coats were removed after germination using forceps, and roots were cut with a scalpel at approximately 5 mm from the root tip. The wound created by the excision was then inoculated with ARqua1 cells. The seedlings were placed on solid Fahräeus medium supplemented with 0.5 mM NH_4_NO_3_ and vertically kept in a growth chamber and weekly transferred into new plates. Transgenic roots were selected based on the fluorescent marker using stereomicroscope (Zeiss, Oberkochem, Germany), and untransformed roots were removed. The composite plants were transferred into pots containing the mixture of sand and vermiculite (1:1, v/v) and kept for one week in a growth chamber before inoculation with rhizobia.

### Confocal laser scanning microscopy

The subcellular localization of DNF2 protein was determined using a TCS SP8 confocal microscope (Leica Microsystems, Wetzlar, Germany) coupled with the LAS X v3 software. For this, transgenic nodules expressing DNF2-eGFP generated via hairy root transformation were harvested and embedded in 6% (w/v) low melting agarose (Biozym Scientific, Germany). Longitudinal sections (70 μm thickness) were made using a VT1000S vibratome (Leica Biosystems, Germany). Sections were then immediately imaged using the confocal microscope with eGFP and mCherry excited at 488 nm and 561 nm, respectively, by White Light Laser (WLL). Emissions were detected with hybrid detectors (HyDs) at 500-550 nm for eGFP, and 600-650 nm for mCherry. Images were analyzed using Fiji/ImageJ software.

### Nodulation assay

*M. truncatula* seeds were germinated in a 22°C incubator for 8-12 h, after which they were grown on nitrogen-free Fahräeus medium plates for 5-7 days. Seedlings were then transferred to pots (6 × 6 × 5.5 cm) containing a sterile 1:1 (v/v) mixture of perlite and vermiculite (four plants per pot). Two days after transplanting, each plant was inoculated with 5 mL of a bacterial suspension adjusted to an OD_600_ of 0.001 (Sm2011 or mutants, as indicated). At 3 and 4 weeks post-inoculation, root systems were harvested, and nodule numbers as well as shoot lengths were recorded.

### Transcriptomics analysis of root nodules

Nodules from *M. truncatula* A17 inoculated with *S. meliloti* mutants (*srp86, srp256, srp485, srp86/srp256, srp86/srp485*) and from *dnf2* (A17 background) inoculated with the wild type strain were harvested at 3 wpi, and total RNA was extracted for paired end sequencing at Novogene (Novogene Co., Ltd., Beijing, China). Raw sequencing reads were processed and gene expression was quantified as described above for *S. meliloti* secretome analysis. Differential expression was conducted using the *edgeR* (v4.0.6) package (*62*), with a unified negative-binomial generalized linear model across all libraries, so that every genotype-to-WT comparison was rigorously adjusted for batch effects within a single fit. Following filtration via *filterByExpr*, TMM normalization was applied. Dispersions were then estimated using *estimateDisp* (robust mode) and integrated into a quasi-likelihood fit with *glmQLFit*. Each mutant was tested against WT with the fold-change–aware quasi-likelihood test glmTreat (|log_2_FC| > 1); genes with Benjamini-Hochberg FDR < 0.05 were considered as differentially expressed.

Genome-wide concordance between *srp86*/*srp485* and *dnf2* was measured as the Pearson correlation of per-gene log_2_(fold changes) (versus WT). A direction-resolved recapitulation fraction was computed for each mutant as the percentage of *dnf2* down– or up-regulated genes that it reproduced with the same sign at FDR < 0.05. GO and KEGG over-representation analyses were run independently on each directional DEG set (background = the 26,762 expressed genes) with clusterProfiler (GO from Blast2GO assignments, tested per ontology; KEGG from KofamScan KEGG-Orthology assignments mapped through the KEGG REST API, excluding global overview maps). Because lineage-specific symbiotic genes (NCR peptides, leghemoglobins, nodule-specific genes) are poorly represented in GO/KEGG (down-set GO coverage 60-61% versus 76% genome-wide), a legume-aware curated enrichment of seven nodule categories was additionally tested by right-tailed Fisher’s exact test against the same background, with Benjamini-Hochberg correction within set.

### Quantification and statistical analysis

Statistical analyses were implemented in GraphPad Prism 7 (GraphPad Software, San Diego, CA). Means were compared using two-tailed Student’s *t* test. Inter-group significance was used one-way ANOVA with Tukey’s test. Samples size (n) and statistical tests are provided in the figure legends.

### Data and code availability

The genetic materials used in this study are available from the corresponding authors upon request. The AlphaFold3 predicted models were publicly accessible at: https://doi.org/10.5281/zenodo.19128292. The code and scripts used in the analysis are publicly deposited at: https://github.com/chongjing/AlphaFold3_Medicago.

## ACKNOWLEDGMENTS

1. P. Liang was supported by the Biological Breeding-National Science and Technology Major Project (Grant No. 2024ZD04079), the Young Scientists Fund (B) of the National Natural Science Foundation of China (Grant No. 32622009), the National Natural Science Foundation of China (Grant Nos. 32470259 and 32300216), the National Key Research and Development Program of China (Grant No. 2024YFA0918200), the Chinese Universities Scientific Fund (2025TC152), and the Pinduoduo-China Agricultural University Research Fund (Grant No. PC2024B02002). G. Zhang and T. Ott were supported by the Enabling Nutrient Symbioses in Agriculture (ENSA) project, which was funded by Gates Agricultural Innovations (G1164932/57461). T. Ott was supported by the Deutsche Forschungsgemeinschaft (DFG, German Research Foundation) under Germany’s Excellence Strategy grant CIBSS-EXC-2189-Project ID 39093984. J.-P. Gao was supported by a Humboldt Research Fellowship from the Alexander von Humboldt Foundation. J.-P. Gao thanks Xiaoyan Shi for support.

## AUTHOR CONTRIBUTIONS

J.-P. Gao and C. Xia conceived the original idea and supervised this project. C. Xia and J.-P. Gao performed AlphaFold3 predictions and RNA-seq analysis. J.-P. Gao and G. Zhang carried out interaction validation. F. Zhao and P. Liang generated rhizobial mutants and performed phenotypic analysis. G. Zhang conducted subcellular localization experiments. Q. Chen contributed to structural prediction and model interpretation. S. Wu performed synteny analysis of DNF2 orthologs. J. Huang designed the website. J.-P. Gao and C. Xia wrote the manuscript, with input from P. Liang, G. Zhang, S. Eves-van den Akker, T. Ott and J. D. Murray. All the authors participated in discussions and data interpretation.

## Supplementary information

**Figure S1.**
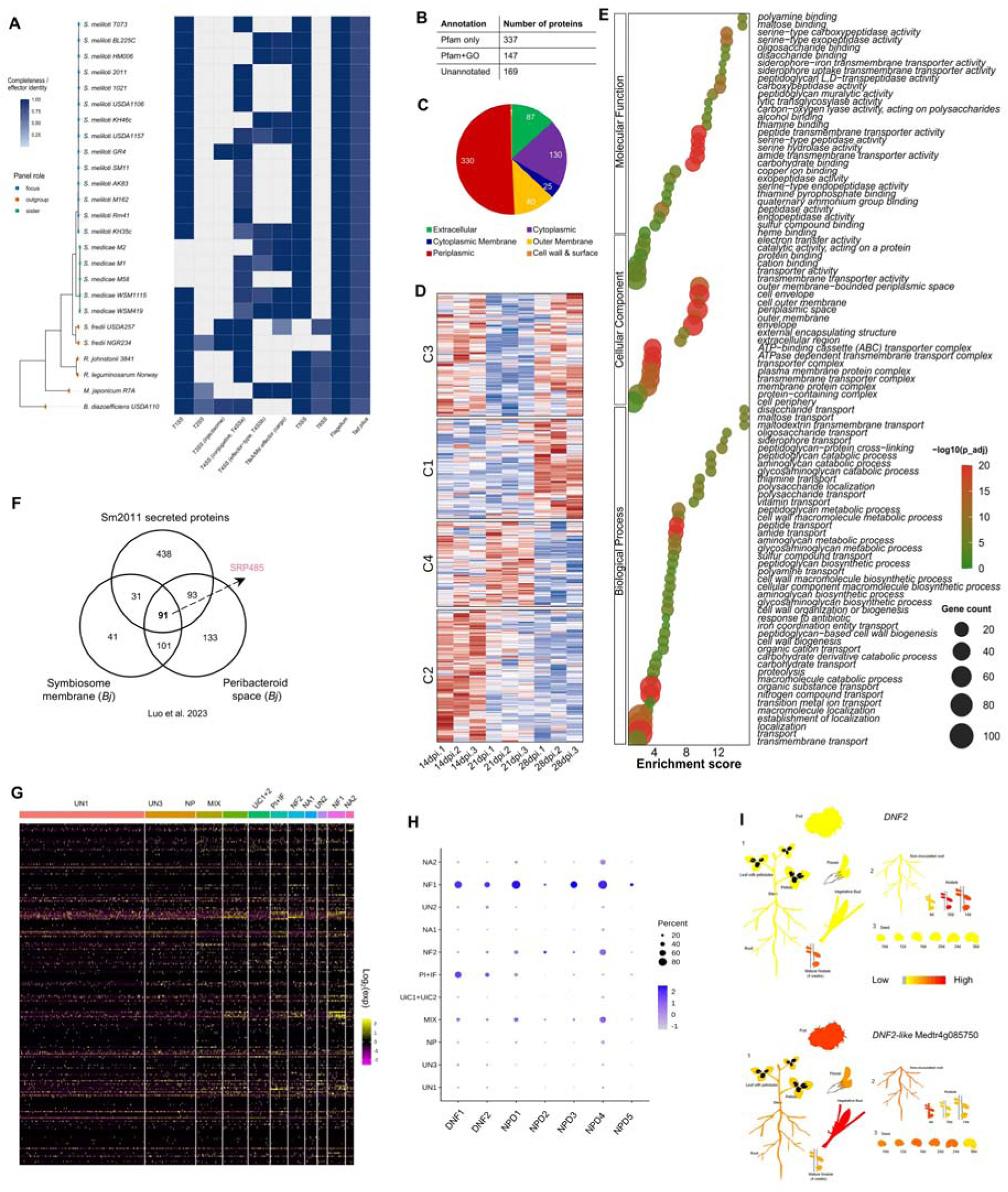
Genomic and transcriptomic characterization of secreted rhizobial proteins and host symbiotic genes during nodule development. (**A**) Analysis of rhizobial secretion systems. Grey indicates not detected. **(B)** Categorization of predicted secreted proteins from *S. meliloti* strain 2011 based on Pfam (http://pfam.sanger.ac.uk/) and Gene Ontology (GO) annotations. **(C)** Predicted subcellular localization profiles of *S. meliloti* proteins. **(D)** Time-course transcriptional expression analysis of *S. meliloti* genes at 14, 21, and 28 days post inoculation (dpi). Data re-analyzed from Sauviac et al. PMID: 35781677. **(E)** GO enrichment analysis of *S. meliloti* genes. **(F)** Venn diagram of *S. meliloti* (Sm) secreted rhizobial proteins and *B. japonicum* (Bj) homologs in Sm. Data re-analyzed from Luo et al. PMID: 35470091. **(G)** An overview of single-cell transcriptome data of 227 nodulation-related genes in Medicago nodules at 14 days post-inoculation. UN, unknown; NP, nodule parenchyma; NF, nitrogen fixation; NA, nodule apex; PI, cell. Data re-analyzed from Ye et al. **(H)** DNF and NPD genes are highly expressed in the infection and nitrogen fixation zones of the nodule. **(I)** Transcript abundance of DNF2 across various Medicago tissues. The data visualization was generated using ePlant (https://bar.utoronto.ca/eplant_medicago/).

**Figure S2.**
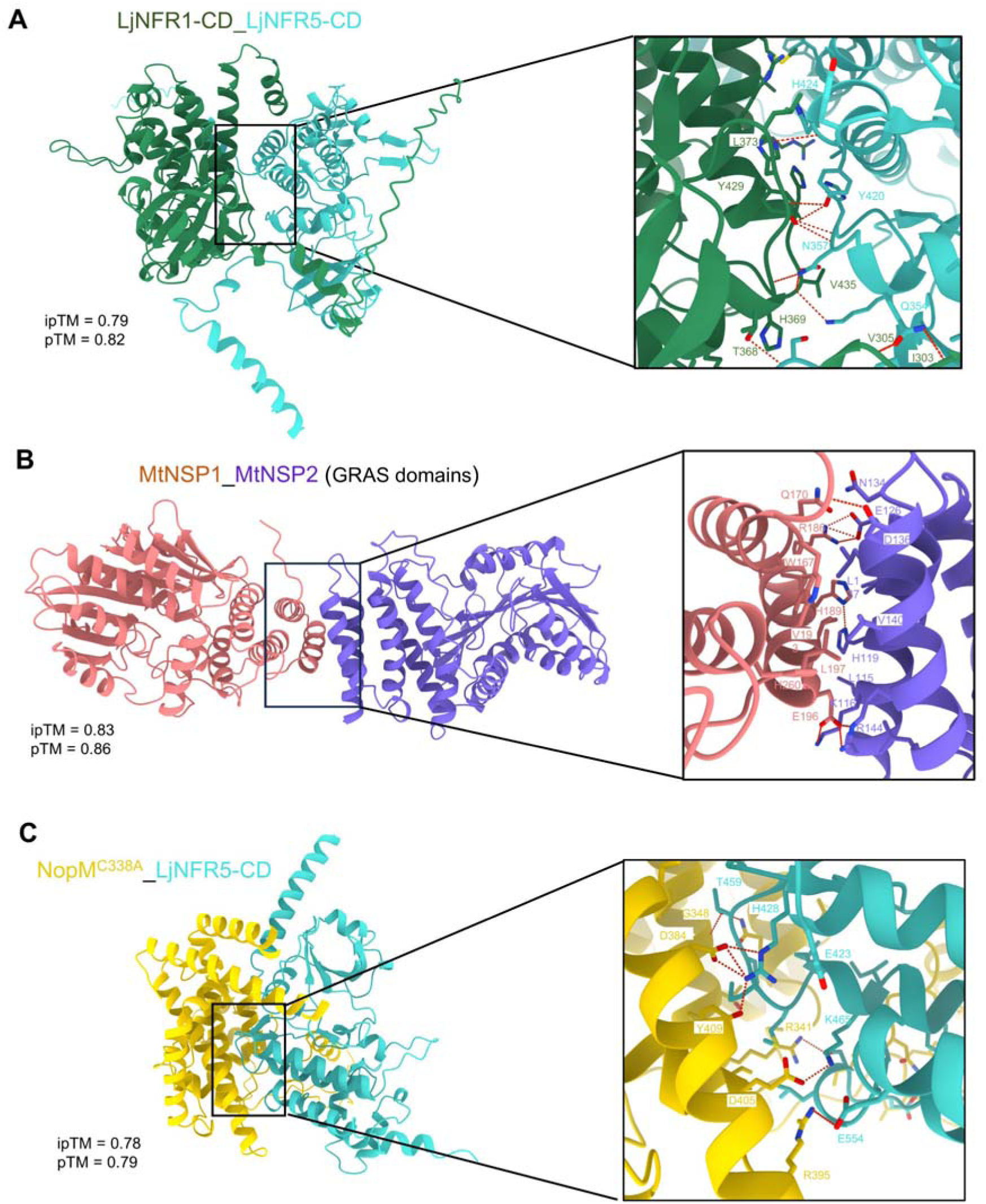
Prediction of known protein complex structures using AlphaFold3. **(A)** Interaction between the intracellular domains of the *Lotus japonicus* Nod factor receptors LjNFR1_CD and LjNFR5_CD. **(B)** Interaction between the *Medicago truncatula* transcription factors NSP1 and NSP2 (GRAS domains). **(C)** Interaction between the T3SS effector NopM and the LjNFR5-CD. The ipTM (interface predicted template modeling) and pTM (predicted template modeling) scores are indicated below each structure. The magnified view shows the interaction interface. Red dashed lines indicate potential hydrogen bonds, and key residues are highlighted.

**Figure S3.**
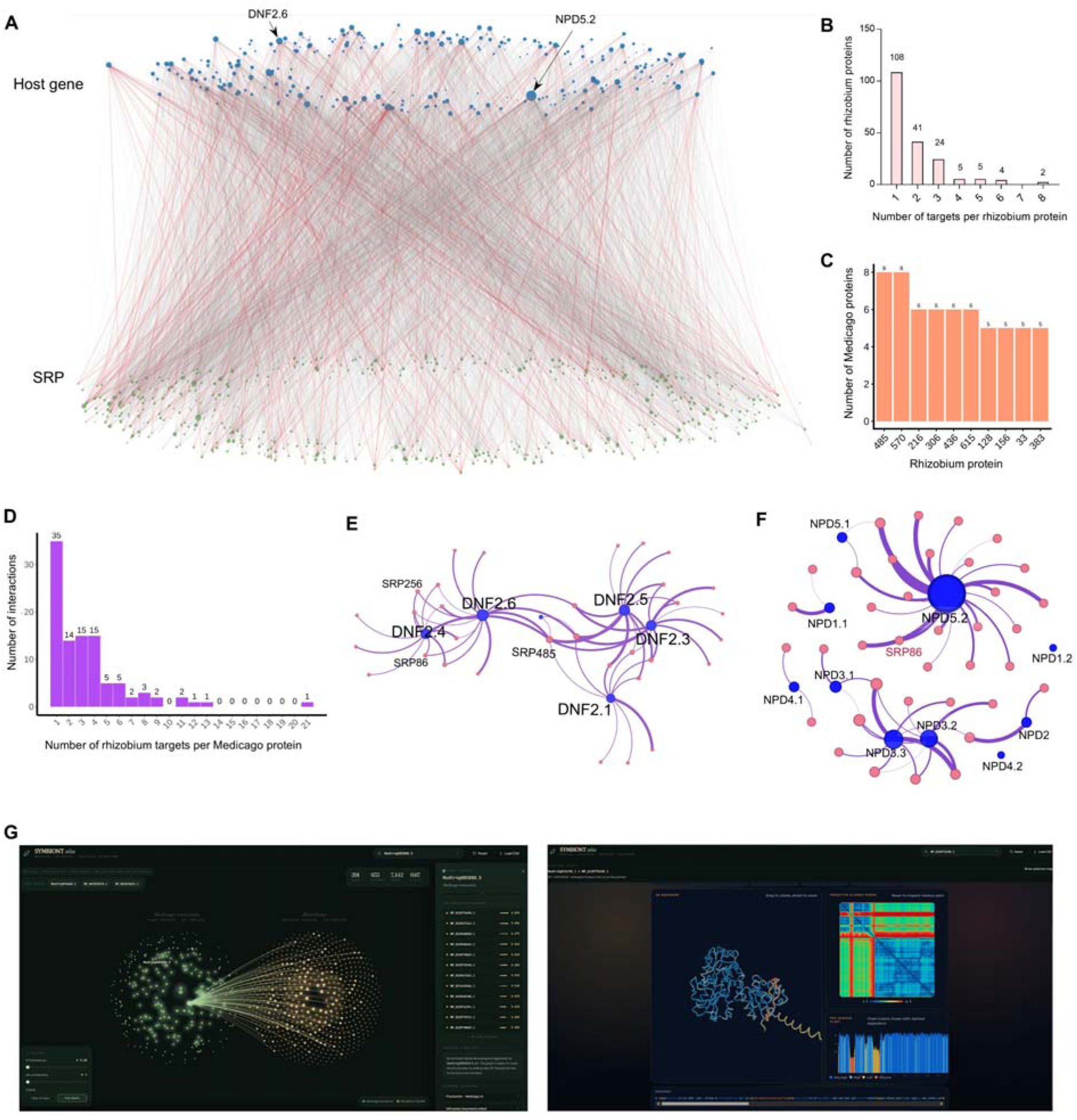
Predicted interactions between rhizobial and plant proteins. **(A)** Interaction network between host genes and secreted rhizobial proteins (SRPs). Red lines represent predicted interaction scores (0.5 × ipTM + 0.5 × pTM) greater than 0.80, while gray lines indicate scores above 0.60. **(B)** Distribution of the number of SRPs interacting with each *Medicago* protein at a score threshold greater than 0.80. 108 SRPs were predicted to interact with a single plant protein, 41 with two different plant proteins, and 24 with three plant proteins. **(C)** The top 10 SRPs with the highest number of interacting *Medicago* proteins, at a confidence score threshold greater than 0.80. SRP485 and SRP570 each interacted with eight distinct plant proteins, representing the highest number among all predicted interactions. **(D)** Distribution of the number of *Medicago* proteins targeted per rhizobial protein at a score threshold greater than 0.80. **(E)** Network visualization of the interactions between DNF2 isoforms and SRPs. Blue and pink nodes represent plant and rhizobial proteins, respectively; node sizes are proportional to the number of interacting partners. Edges denote predicted interactions with scores above 0.80. **(F)** Network visualization of the interactions between NPD and SRPs. **(G)** An example screenshot of the legume-rhizobia interactome webpage.

**Figure S4.**
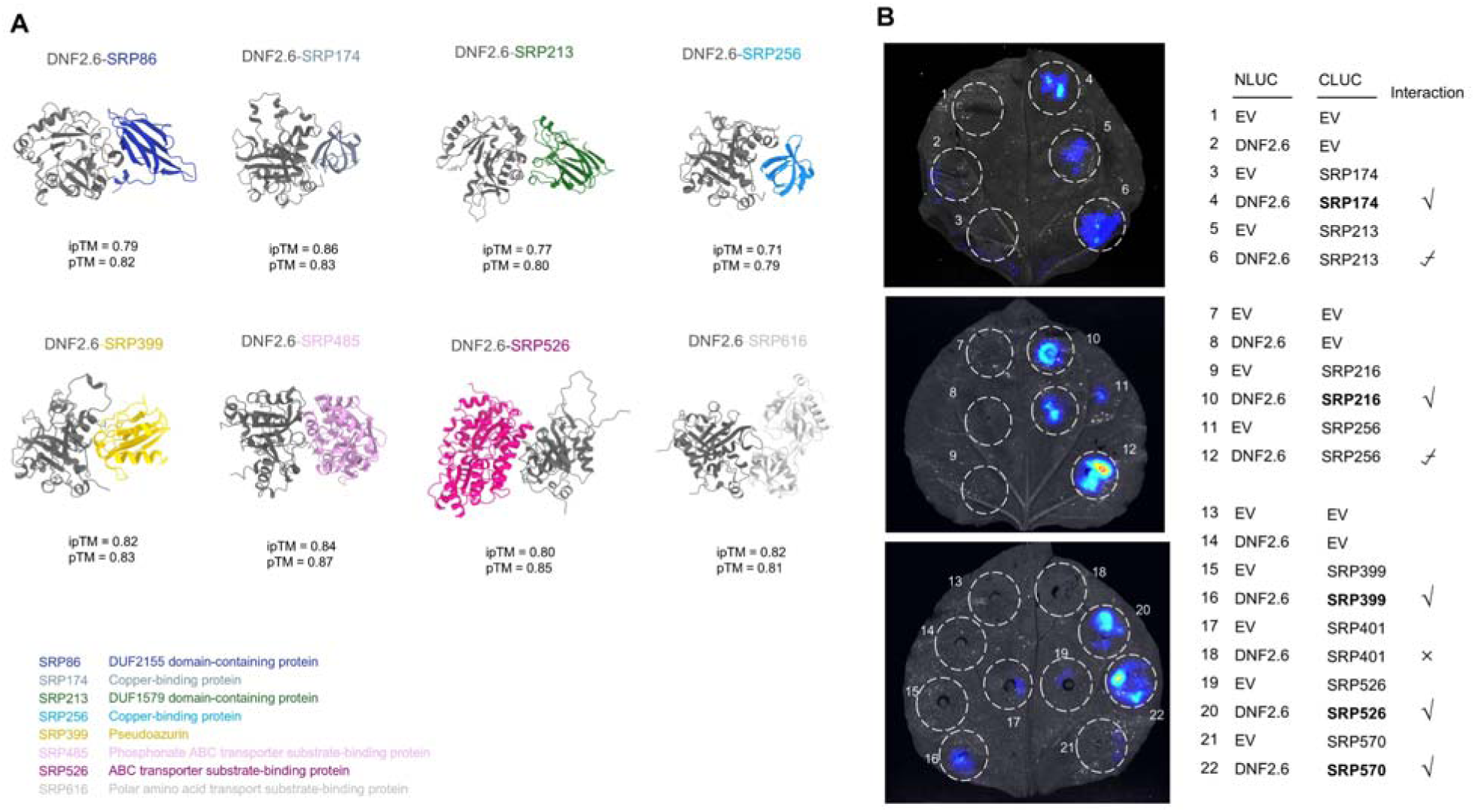
DNF2 interacts with SRPs. **(A)** The predicted interfaces of DNF2.6 with different SRPs. The SRPs are color-coded for clarity. The ipTM (interface predicted template modeling) and pTM (predicted template modeling) scores are indicated below each structure. **(B)** Split-luciferase complementation assays validating the interaction between DNF2 and various SRPs in *Nicotiana benthamiana*. Luminescence signal intensity correlates with binding strength, with indicated by strong (⍰), weak (⍰), or no interaction (×).

**Figure S5.**
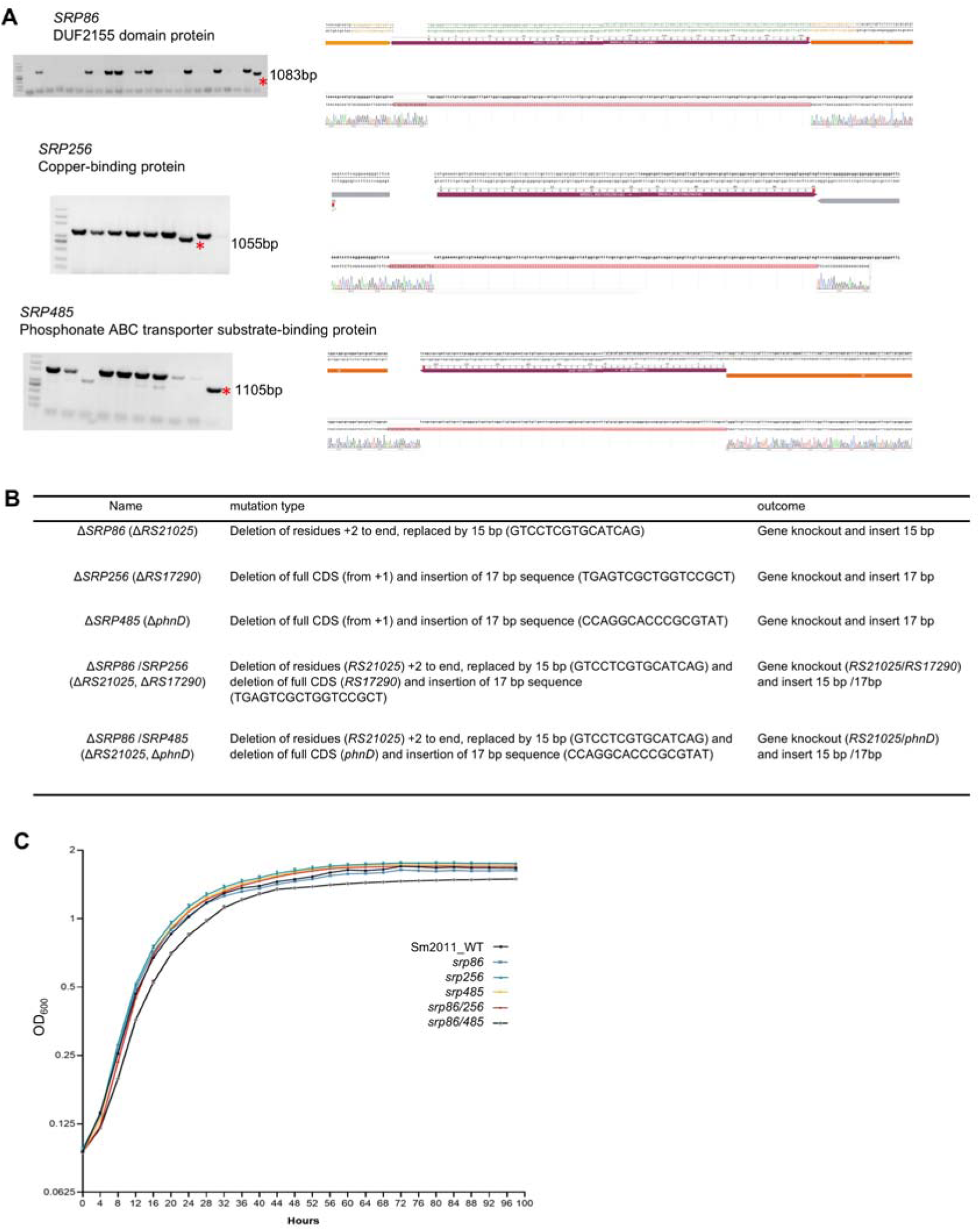
Construction of Rhizobium mutants. **(A)** Identification of rhizobia mutants. Left panel: agarose gel electrophoresis; red asterisks indicate bands corresponding to knockout mutants. Right panel: sequencing chromatograms. **(B)** Mutation profile of the *srp* mutants. Detailed genomic insertion and/or deletion patterns displaying the precise nucleotide alterations within the *srp* locus compared to the wild-type strain. **(C)** Comparative growth kinetics of rhizobia strains. Dynamic growth curves of wild-type and mutant strains cultured in liquid tryptone-yeast extract medium. Optical density (OD_600_) was recorded at 4 h intervals over a total period of 100 h.

**Figure S6.**
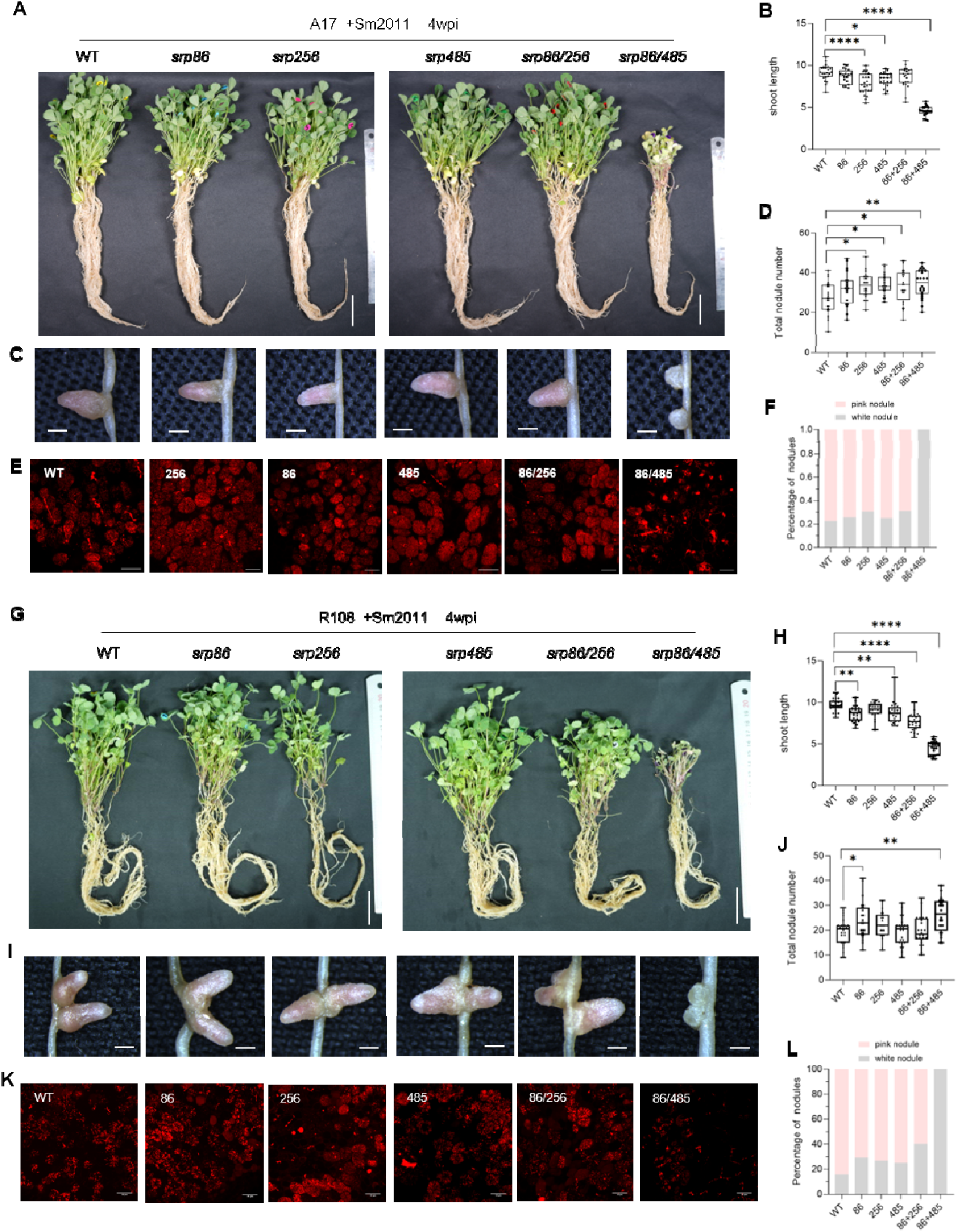
Plant growth and nodulation phenotypes 4 weeks post inoculation with the *srp* strain. (**A, G)** Growth phenotypes of *Medicago truncatula* A17 (A) or R108 (G) with either the wild-type (WT) rhizobial strain Sm2011 or the indicated mutant rhizobia. Photographs were taken 4 weeks post-inoculation (wpi). Scale bar = 3 cm. **(B, H)** Quantification of shoot length in genotype A17 (B) or R108 (H). Boxes show the first quartile, median, and third quartile, whiskers show minimum and maximum values, dots show data points (n = 24). **(C, I)** Representative stereomicroscopy images of mature nodules harvested at 4 wpi. Scale bars = 500 µm. **(D, J)** Total nodule numbers per plant in genotype A17 (D) or R108 (J) at 4 wpi. **(E, K)** Confocal microscopy images of nodule nitrogen-fixation zones. Red fluorescence indicated Sm2011 expressing mCherry. Scale bars = 50 µm **(F, L)** Quantification of the proportion of pink versus white nodules. The above experiments were independently repeated three times with similar results. Statistical analysis was performed using one-way ANOVA followed by Tukey’s HSD test (*, *P*<0.05, ** *P*<0.01, ****, *P*<0.0001).

**Figure S7.**
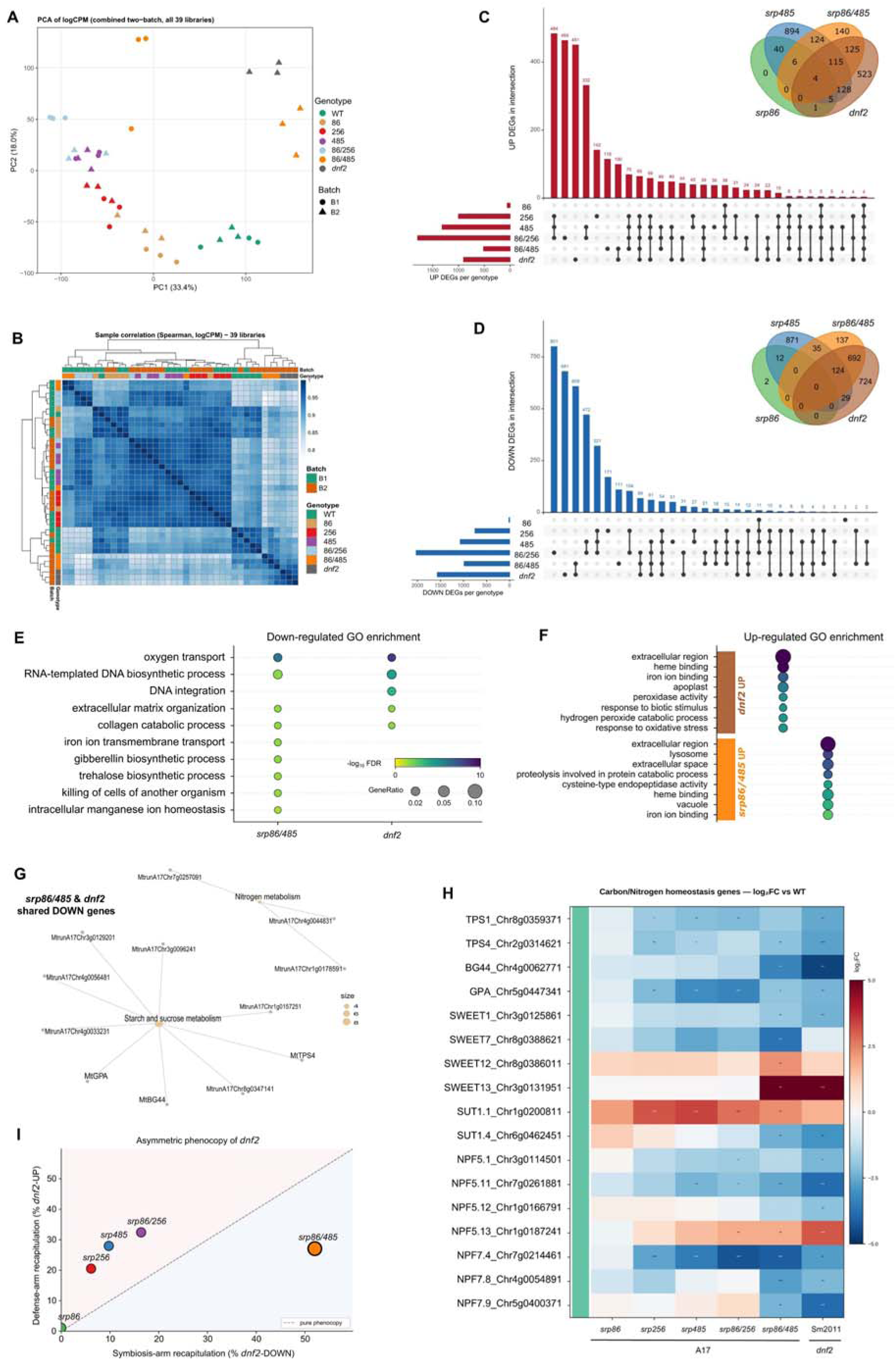
Transcriptomic analyses of host and bacterial mutants. **(A)** Principal component analysis (PCA) of global expression profiles across genotypes. **(B)** Sample-to-sample correlation. Heatmap of pairwise Spearman’s rank correlation coefficients computed on logCPM values across all filtered genes for the 21 libraries, hierarchically clustered (Euclidean distance, complete linkage). **(C, D)** Intersection of differentially expressed genes (DEGs) across genotypes. UpSet plots of genes significantly up-regulated (C) and down-regulated (D) versus WT (glmTreat, |log₂FC| > 1, FDR < 0.05) in each of the six mutants. The Venn diagram shows the overlap of differentially expressed genes among *srp86*, *srp485*, *srp86/485*, and *dnf2* (all compared to wild-type nodules), with down– and up-regulated genes presented separately. **(E)** Top Gene Ontology (GO) biological-process terms enriched among down-regulated genes *srp86/485* versus *dnf2*); both converge on oxygen transport and related nitrogen-fixation/oxygen-buffering processes. **(F)** Divergence of top GO terms of up-regulated genes. **(G)** KEGG pathway gene-concept network of the shared down-regulated signature between *srp86/485* and *dnf2* vs wild-type. **(H)** Direction-resolved recapitulation of the *dnf2* signature by each bacterial mutant. Percentage of *dnf2* down-versus up-regulated genes reproduced from each mutant was shown in x– and y-axis, respectively.

**Figure S8.**
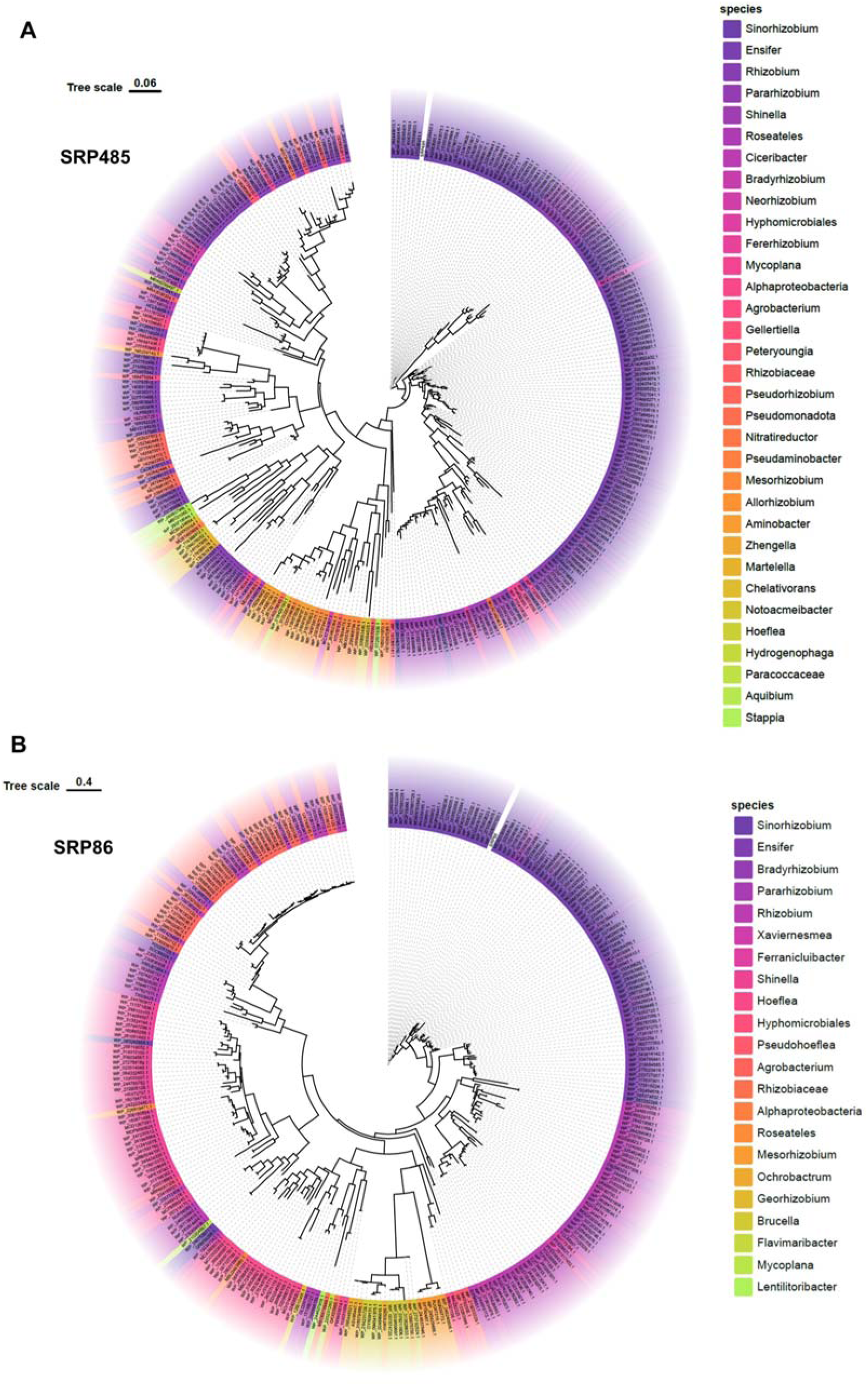
Phylogenetic trees of SRP485 and SRP86 homologs across representative rhizobial strains. **(A, B)** The tree includes genera such as *Sinorhizobium*, *Ensifer*, *Bradyrhizobium*, *Rhizobium*, and others. Homologous sequences were retrieved from the NCBI nr database (E-value ≤ 1 × 10^-5^, sequence identity ≥ 80%). For each seed sequence, the top 300 hits with the highest bit-scores were selected and combined with the corresponding seed sequence (301 sequences per seed). Multiple sequence alignment was performed using MAFFT, and phylogenetic trees were constructed with FastTree. Branch lengths represent evolutionary distances.

**Table S1.** Profile of *Sinorhizobium* and *Medicago* genes analyzed in this study.

**Table S2.** Predicted cross-kingdom interactions identified by AlphaFold3.

**Table S3.** Comparative transcriptomic profiling of nodules formed by *srp* rhizobial mutants or on *dnf2* mutant host plants.

**Table S4.**
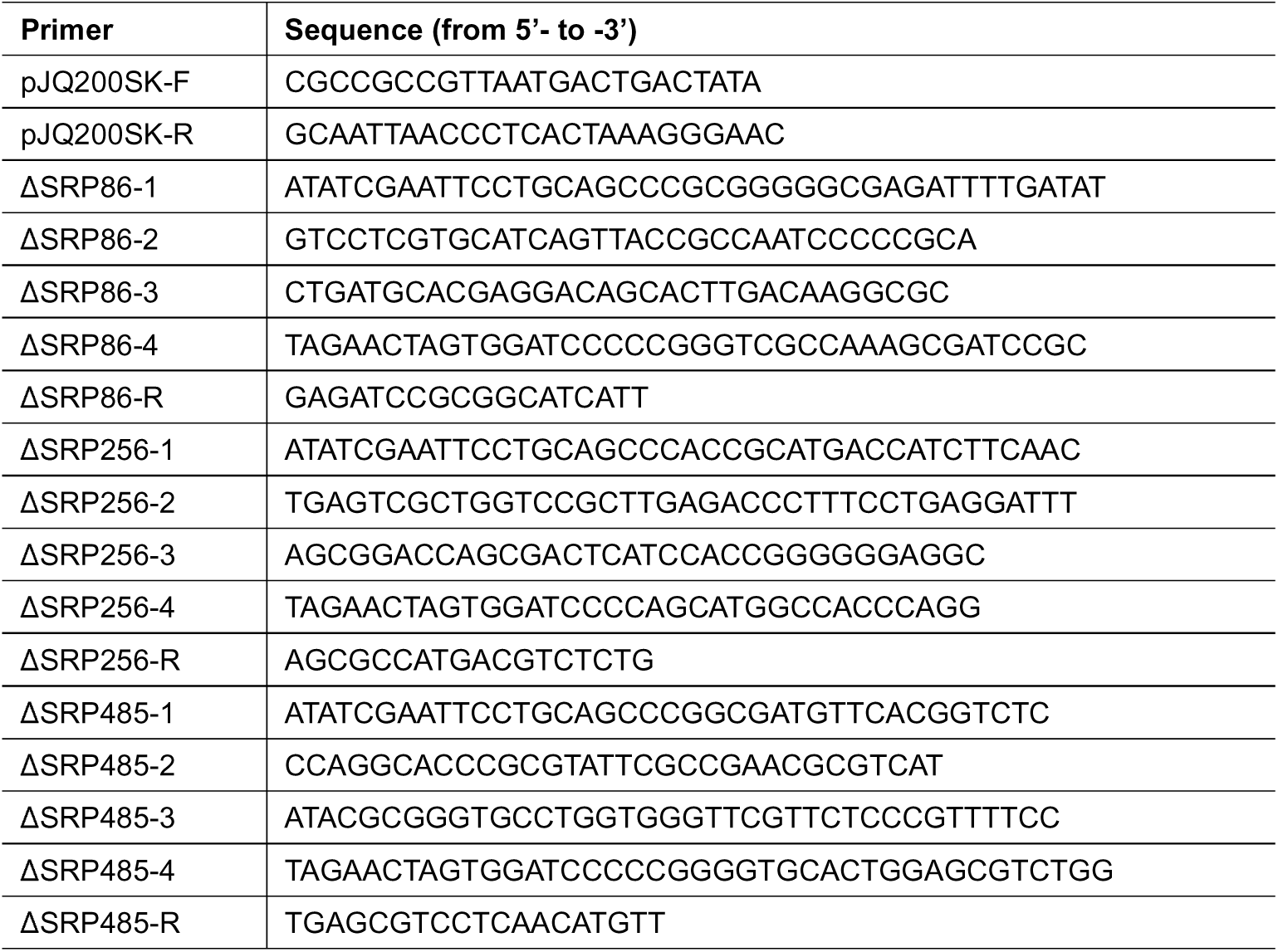
Primers used in this study.

